# Tractography-based transcranial magnetic stimulation prediction using machine learning

**DOI:** 10.1101/2025.11.11.685674

**Authors:** Lucas dos Santos Betioli, Renan Hiroshi Matsuda, Thais Cunha Marchetti, Carlo Rondinoni, Victor Hugo Moraes, Márcio Adriano de Campos Júnior, Carlos Ernesto Garrido Salmon, Renato Tinós, Oswaldo Baffa Filho

**Affiliations:** Department of Physics, Faculty of Philosophy, Sciences and Letters of Ribeirão Preto (FFCLRP), University of São Paulo (USP), Av. Bandeirantes, 3900, Monte Alegre, 14040-901, Ribeirão Preto, SP, Brazil; Department of Neuroscience and Biomedical Engineering, Aalto University School of Science, Rakentajanaukio 2, 02150, Espoo, Finland; Department of Medical Imaging, Hematology and Clinical Oncology, Ribeirão Preto Medical School (FMRP), University of São Paulo (USP) and Ribeirão Preto Medical School Hospital das Clínicas, Av. Bandeirantes, 3900, 14049-900, Ribeirão Preto, SP, Brazil; Department of Computing and Mathematics, Faculty of Philosophy, Sciences and Letters of Ribeirão Preto (FFCLRP), University of São Paulo (USP), Av. Bandeirantes, 3900, Monte Alegre, 14040-901, Ribeirão Preto, SP, Brazil

**Author notes:** Corresponding author: Lucas dos Santos Betioli, E-mail(s). Equal contribution in first authorship (Lucas dos Santos Betioli, Renan Hiroshi Matsuda).

**Keywords:** Transcranial Magnetic Stimulation, Machine Learning, Tractography, Responsivity, Motor Threshold, Convolucional Neural Networks, Motor Mapping

## Abstract

**Background:** Identifying the stimulation brain target is fundamental for transcranial magnetic stimulation (TMS). Currently, this process is time-consuming and heavily dependent on the operator’s expertise.

**Objective:** This study evaluated a deep learning–based approach to structural brain connectivity for improving stimulation site prediction and real-time cortical excitability mapping during neuronavigation.

**Results:** Tractography-derived connectivity features and distal myographic responses were used to train four neural network models across five subjects. Neural network inputs were incrementally varied using either tractography alone, coil coordinates alone, or hybrid combinations of both using different concatenation regimes. The best performance was observed in two subjects out of 5, where hybrid models integrating coil coordinates and fiber information achieved higher F1 scores and accuracy.

**Conclusion:** Despite these promising results, substantial inter-subject variability was observed. This approach may also be applied to other brain regions to investigate connectivity, and the use of machine learning could be extended to functional domains beyond the motor cortex, including sensory and cognitive areas.

## Introduction

Transcranial magnetic stimulation (TMS) is a non-invasive neuromodulation technique that utilizes focused magnetic pulses to modulate neuronal activity in specific regions of the brain (1). In this procedure, a coil is placed externally on the scalp and an electric current is applied. The rapid change in the current generates a magnetic pulse lasting several hundreds of microseconds, which in turn induces electric fields in the brain tissue. These induced electric fields are capable of depolarizing cortical neurons and generating action potentials (2).

In the case of stimulation over the primary motor cortex, a portion of the resulting action potential travels along descending pathways, such as the corticospinal tract, reaching spinal motor neurons and ultimately producing motor responses in peripheral muscles (3). The responses can be measured by an electromyograph using surface electrodes, and this response is referred to as the motor evoked potential (MEP) (4).

A commonly used application of TMS is motor mapping, aiming to investigate the motor cortical representation in humans, most often in hand muscles (5). An important parameter in this procedure is the hotspot, defined as the cortical region directly beneath the center of the coil that produces a MEP at the lowest possible stimulation intensity — known as the resting motor threshold (MT). This quantity is defined as the stimulus intensity capable of evoking peak-to-peak MEP amplitudes larger than 50 µV in at least 5 out of 10 trials (2). This technique gains precision and stability when conjugated with Neuronavigation. The fine tridimensional control of the stimulation coil provides high real-time accuracy, allowing the operator to target patient-specific cortical regions. In its absence, hotspot hunting becomes a time-consuming process dependent on operator’s manual control stability and strength, leading to larger variability in results compared to using a neuronavigation system (6).

Beyond the anatomic magnetic resonance imaging (MRI), the integration of other neuroimaging modalities also increases assertiveness in motor mapping and muscle targeting. Diffusion-weighted images serve as a starting point to reconstruct sub-cortical tracts, in a technique called tractography. The procedure is used to reconstruct neural pathways in the brain ,which the TMS-induced signals propagate and are composed of organized bundles of white matter fibers (7,8).

The current gold standard for target localization depends on the operator skill in manipulating the coil and identifying the motor hotspot (9,10). This paper proposes a Machine Learning (ML) approach based on tractography information to automate the determination of motor cortex targets, optimizing hotspot hunting. Therefore, we propose automating hotspot localization by training and comparing ML models using tractography structures as input and highly reactive point locations as output.

## Materials and Methods

### Participants

Five young adult participants (age: 32 ± 11 years; one female) with no history of neurological disorders took part in this study. All participants provided informed consent, and any discomfort during the experiments was reported. The experimental protocol was approved by the institutional Ethics Committee (approval number: 85778925.0.0000.5407). Participants’ handedness was determined using a modified Edinburgh Handedness Inventory (11), administered online via the Brain Mapping website (https://www.brainmapping.org/shared/Edinburgh.php).

The following sections outline the procedures involved in MRI acquisition and preprocessing, including anatomical segmentation and tractography; the collection and processing of EMG signals to map motor cortex activity. and the creation of a dataset for training and testing the machine learning models. Additionally, we detail the use of several machine learning approaches to predict motor cortex responsiveness based on experimental data.

### MRI acquisition and imaging preprocessing

The T1-weighted structural MRI volume (512 × 512 × 257 voxels; 0.5 × 0.5 × 0.7 mm³ resolution) and diffusion-weighted image (DWI) with an isotropic voxel (2mm³), using a multi-shell protocol with 96 diffusion gradient directions and b-values of 0, 900, 1600, and 2500 s/mm², were acquired on 3T Philips Achieva dStream scanner.

Cortical and subcortical structures were segmented using FreeSurfer’s recon-all pipeline with default parameters, resulting in aparc+aseg parcellation files (12–21). As illustrated in Figure 1, from these parcellations the precentral gyrus was specifically selected to aid in neuronavigation.

**Fig 1.**
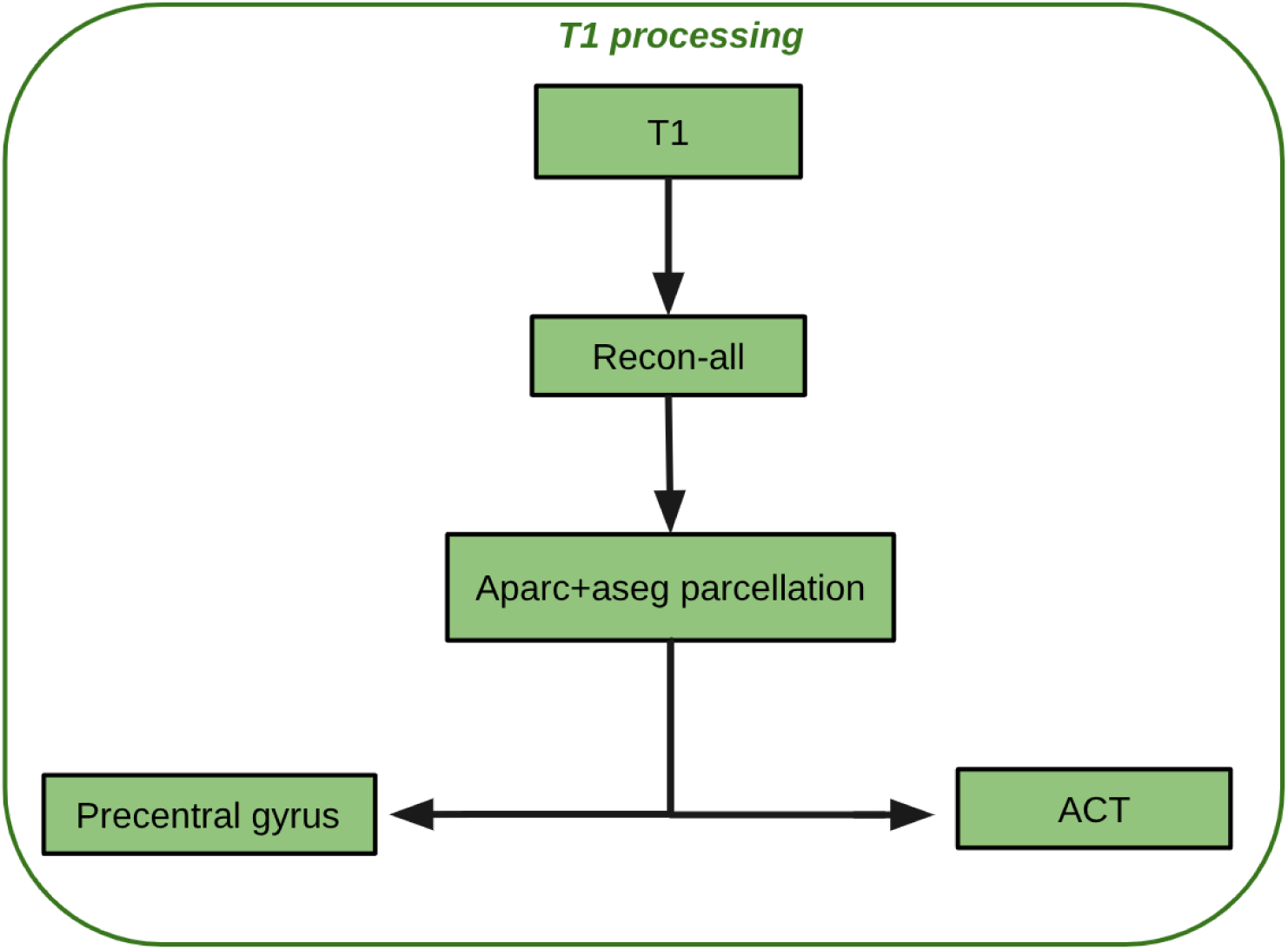
Processing of T1-weighted images and extraction of brain regions. T1-weighted : images were processed using the recon-all pipeline to generate aparc+aseg parcellation files. From these parcellations, the precentral gyrus was specifically extracted for neuronavigation, and other regions were obtained for ACT

The DWI processing pipeline, illustrated in Figure 2, followed the minimal pipeline defined in the previous study (22) and using MRtrix3 software (23). First, a brain mask was created to segment the brain from the surrounding head tissue and to serve as a mask on diffusion volumes. Second, the Marchenko-Pastur principal component (MP-PCA) (24–26) was applied, followed by denoising, and Gibbs-ringing artifact removal (27). Susceptibility distortions, eddy currents, and motion artifacts were jointly corrected using FSL’s standard topup/eddy workflow with reverse-phase encoded B₀ pairs. (28–31). Finally, bias-field correction was performed using FSL (29,32).

**Fig 2.**
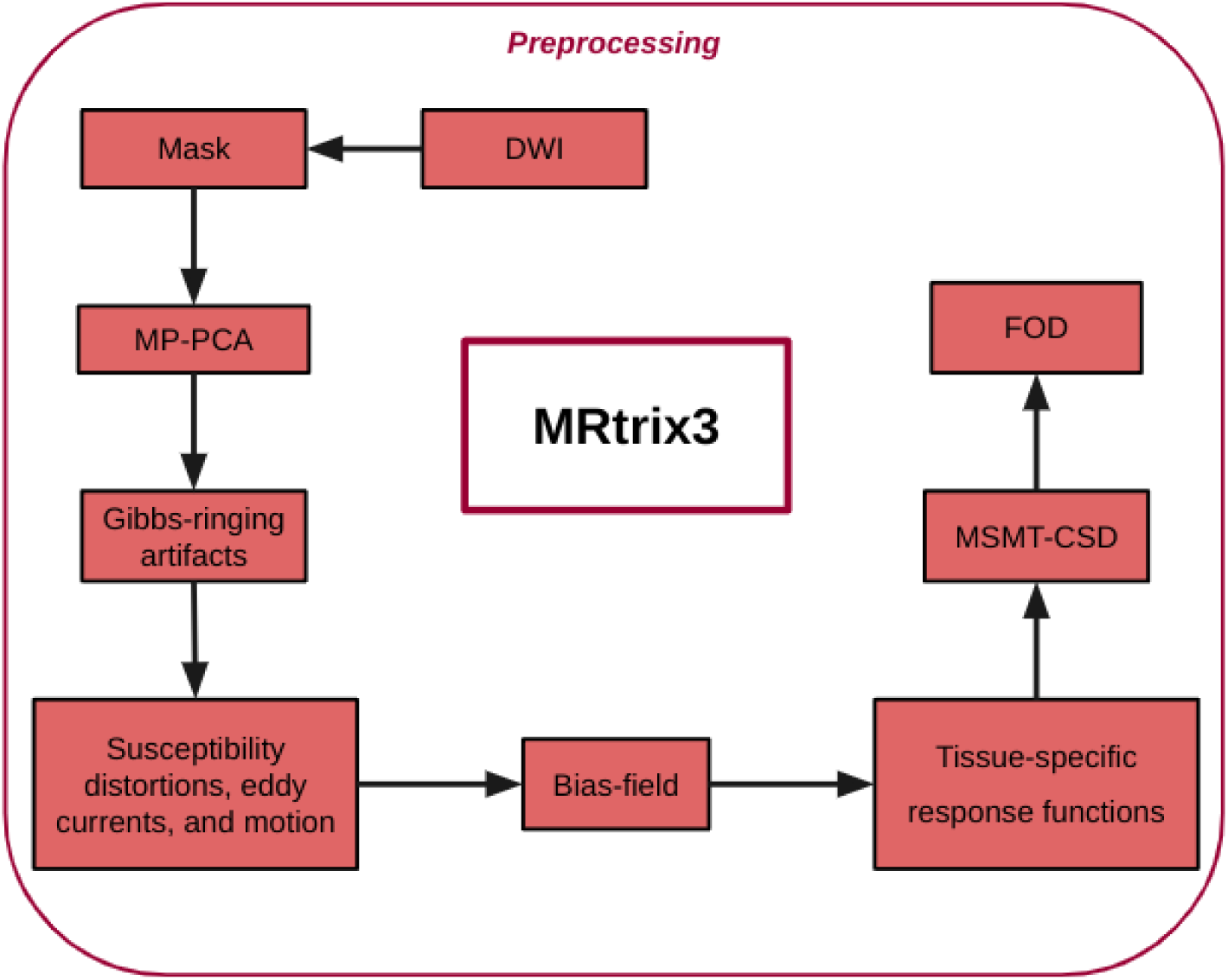
Preprocessing pipeline for DWI using MRtrix3: The workflow includes brain masking, MP-PCA denoising, Gibbs-ringing artifact removal, correction for susceptibility distortions, eddy currents, and motion, and bias field correction. Tissue-specific response functions were estimated using the Dhollander method, followed by MSMT-CSD to compute the FOD.

**Fig 3.**
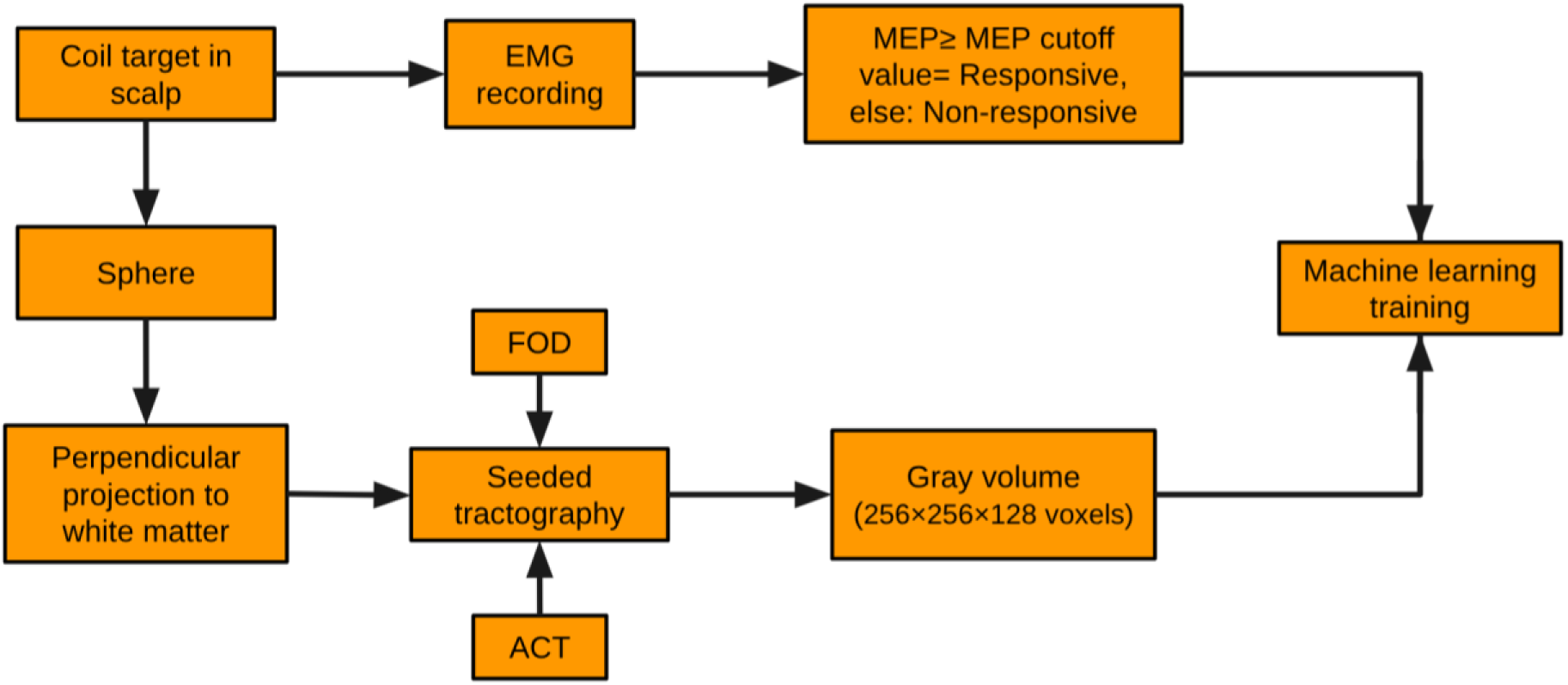
Workflow for tractography-based data processing: For each coil target on the scalp, the coil position was projected onto the white matter surface to define a tractography seed point. Seeded tractography (with ACT constraints) was performed, and the resulting streamlines were converted to grayscale volumes (256×256×128 voxels). In parallel, MEPs were used to classify each target as responsive or non-responsive based on the MEP amplitude cutoff. These labeled volumes were then used to train machine learning models. Detailed procedures are described in the Supplementary Material (subchapter Projection of Stimulation Targets onto the White Matter Surface).

Tissue-specific response functions were estimated via the Dhollander unsupervised method (33,34). Fiber orientation distributions (FOD) were computed using the multi-shell, multi-tissue constrained spherical deconvolution (MSMT-CSD) approach (35). The resulting FOD were subsequently used as input data for the Trekker algorithm, implemented within the InVesalius software, to perform fiber tracking and to visualize the reconstructed fibers.

### EMG acquisition and processing

The motor cortex was initially mapped using TMS in all the volunteers. Stimulation was delivered with a figure-8 coil (C-B70) connected to a MagPro X100 stimulator (MagVenture) for all participants, except for Subject 2, who was stimulated using a MagPro 30 device (MagVenture). In hotspot hunting, the motor threshold was determined by identifying the minimal stimulus intensity capable of reliably eliciting MEPs with amplitudes of at least 50 µV in at least 3 out of 6 consecutive trials at rest (36).

Subsequently, a motor mapping was conducted using stimulation intensities set at 110% of the previously established motor threshold. Approximately 50 target locations per volunteer were stimulated across the motor cortex, with 10 biphasic waveform pulses administered at each location, at randomized intervals ranging from 4 to 6 seconds.

The primary objective of the motor mapping was to stimulate the motor cortex to identify a core responsive region (the hotspot) and its surrounding areas with lower or absent motor responses. Mapping was performed using the hotspot as the central reference point, around which a spatial grid was manually defined. Stimulation sites were defined along lateral and medial directions based on this reference grid, enabling comprehensive coverage of the primary motor region and assessment of the spatial extent of motor responsiveness. At each stimulation site, the coil angles and coordinates were recorded, and the corresponding mean motor-evoked potential amplitude was calculated for each location. Coil localization and positioning were executed using a Dobot CR10 robotic arm, tracked by the Polaris Vega VT system. Both devices are controlled by the InVesalius Neuronavigator software (37–39) with a robotic positioning threshold of 2 mm and 2°. The coil was positioned perpendicular to the motor cortex to maximize the motor response detected by EMG.

A robotic arm (left) holds the TMS coil in a stable and reproducible position over the subject’s scalp. Reflective markers attached to both the coil and the subject’s head enables real-time position tracking via an optical tracking device (upper right). This tracking system communicates with the neuronavigation software InVesalius Neuronavigator, which continuously registers the coil position relative to the subject anatomy and predefined stimulation targets. The subject remains seated with surface electromyography (EMG) electrodes placed on the right hand muscles to record motor evoked potentials (MEPs).

EMG signals of the MEP were recorded in all subjects. Specifically, data from volunteers 2 and 4 were collected using the Neurosoft Neuro-MEP-Micro device at a sampling frequency of 20 kHz, whereas data from the other volunteers were recorded using the Bittium NeurOnE system at 5 kHz per channel. To ensure data consistency across acquisition systems, signal stability was verified by confirming that the signal-to-noise ratio (SNR) remained within a comparable range for both devices and that MEP amplitudes fell within the expected physiological range across sessions. Electrodes were placed following the belly-tendon montage in the flexor pollicis brevis. The signal was preprocessed with an eighth-order, high-pass Butterworth filter with a 20 Hz cutoff frequency and notch filtering at 60, 120, 180, 240, 300, and 360 Hz to minimize powerline interferences.

For each target, the MEPs were recorded over 10 stimuli and the corresponding coil target coordinates were collected. Using this data, the Training Dataset was generated, where each motor mapping target was transformed into a seed point for tractography. This procedure is described in detail below.

### Training Dataset

The dataset used for training the machine learning models was composed of post-processed fiber tracts obtained with Trekker, implemented within the InVesalius software. For tractography, each stimulation site recorded by the neuronavigation system was transformed into a seed point, from which fiber tracking was performed to visualize neural pathways. Subsequently, the tractography results were reconstructed into standardized volumetric maps and converted into grayscale images. A description of the seed-point definition and tractography procedures is presented below.

For each stimulation location, EMG signals were recorded, and the coil orientation and position were represented as a single transformation matrix, combining Euler angles (α, β, γ) and Cartesian coordinates (X, Y, Z). Each stimulation location was then projected perpendicularly onto the white matter surface, and this projection defined the seed point for tractography

Tractography was then performed in the InVesalius using the Real-Time module with Trekker’s default parameters (seed radius = 1.5 mm; maximum 500 streamlines per seed) (40). To capture probabilistic variability, each seed was processed five times, generating multiple tractograms per subject. Each tractogram was treated as augmented data for training the machine learning models. These tractograms were exported containing coordinates (x, y, z), RGB color values, and opacity.

The parcellations obtained from aparc+aseg files (41) were subsequently used to make an ACT using the Trekker ACT script (42), which restricts streamline generation to biologically plausible pathways, effectively pruning spurious fibers and limiting false-positive streamline(40).

A custom Python pipeline reconstructed volumetric maps from the tractograms. First, a downsampled T₁-weighted volume (256×256×128 voxels), derived from the original 512×512×257 dataset, provided an empty canvas for voxel insertion. For each tractogram, the script parsed world-space coordinates, RGB intensities, and opacity values. Voxel values were converted from RGB to a single grayscale channel using the luminosity formula (0.299 R + 0.587 G + 0.114 B), multiplied by the corresponding opacity to weight the intensity. After, coordinates were translated into voxel indices via the inverse NIfTI affine and integer casting, voxel values were converted to grayscale.

In parallel with this pipeline, median MEP amplitudes were calculated and used to classify each tractography volume as either *responsive* or *non-responsive*. Using the median MEP amplitude as a threshold ensured a balanced distribution between responsive and non-responsive classes.

### Machine learning models

For each subject, four distinct classification models were evaluated. Each model was trained with its own set of hyperparameters. All four models trained for predicting cortical responsiveness used the Adam optimizer for parameter optimization (Kingma and Ba 2014). Regarding the learning rate, Model 1 (Coordinates + Angles) employed a rate of 0.0001, whereas Models 2 (Fibers Only), 3 (Fibers + Coordinates V1), and 4 (Fibers + Coordinates V2) used a lower learning rate of 0.000001. In terms of training epochs, Model 1 was trained for 300 epochs, while the other three models were trained for 30 epochs each.

This specific configuration of hyperparameters reflects differences in model complexity and convergence behavior. All models employed one-hot encoded labels and a softmax output layer for binary classification. The output consisted of a two-dimensional vector, where each component corresponds to one class (non-responsive or responsive). Although these values are not strict probabilities in the statistical sense, the softmax function ensures that they sum to one, allowing them to be interpreted as relative likelihoods assigned by the model to each class. An overview of the models used in this study is presented in Figure 4, and their specific architectures and input features are described below:

- Model 1 (MLP): A multilayer perceptron trained solely on the coil’s Cartesian coordinates and Euler angles. This model tests the hypothesis that it is possible to predict the hotspot using only the coil position and orientation, without any fiber information.
- Model 2 (LeNet-5): A convolutional neural network inspired by the LeNet-5 architecture, trained exclusively on 257 × 257 × 126 voxel tractography volumes. This model investigates whether the tractography data alone contains sufficient information to predict the hotspot, independent of coil coordinates and angles.
- Model 3 (V1 hybrid): An early-fusion dual-input model where the coil parameters are concatenated with the flattened convolutional features in the first fully connected layer. This model is designed to evaluate whether combining both coil parameters and tractography features leads to improved hotspot prediction compared to using each modality separately.
- Model 4 (V2 hybrid): A late-fusion architecture in which coil parameters are processed through an MLP branch and tractography volumes through a convolutional branch, with both paths merging at the final classification layer. This model tests whether processing both data types independently and fusing them only at the classification stage (late fusion) outperforms the early-fusion approach of Model 3.

**Fig 4.**
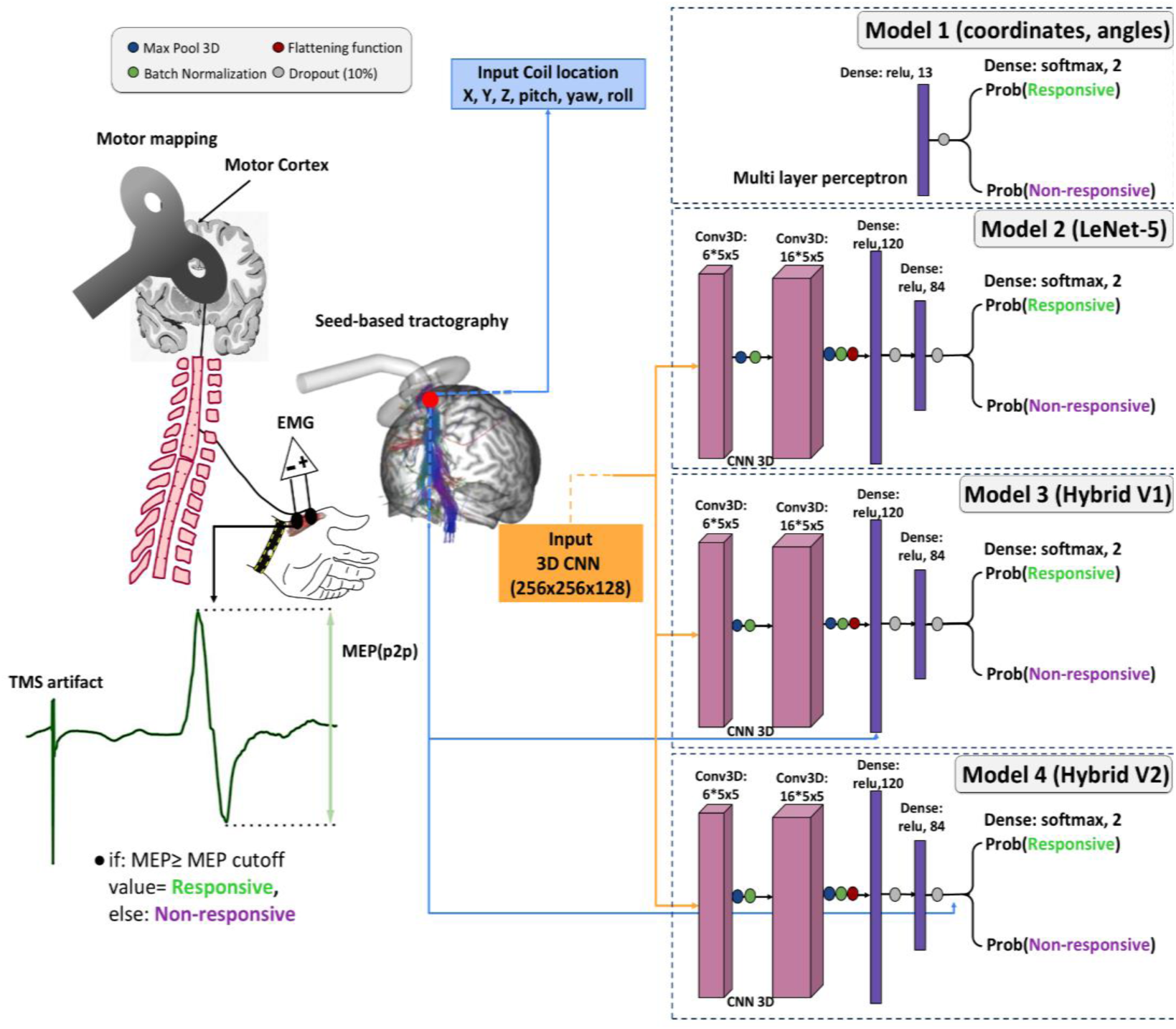
Workflow for tractography-based model training: Motor mapping is performed through EMG recordings of MEP peak-to-peak amplitudes elicited by stimulation of the motor cortex, with sites classified as responsive or non-responsive depending on whether the mean MEP amplitude exceeds the cutoff threshold. Coil position and orientation parameters (X, Y, Z, pitch, yaw, roll) are used as inputs for Model 1 (multilayer perceptron), while volumetric brain images (256×256×128) serve as inputs for the 3D CNN architectures (Models 2–4). Model 2 follows a LeNet-5–based CNN, whereas Models 3 and 4 represent hybrid architectures combining CNN feature extraction with dense layers of increasing depth (Hybrid V1 and V2, respectively). Each model outputs the probability of responsiveness (Prob[Responsive]) and non-responsiveness (Prob[Non-responsive]). This predictive module is designed to be integrated into InVesalius Navigator, allowing real-time responsivity prediction during motor mapping procedure.

**Fig 5.**
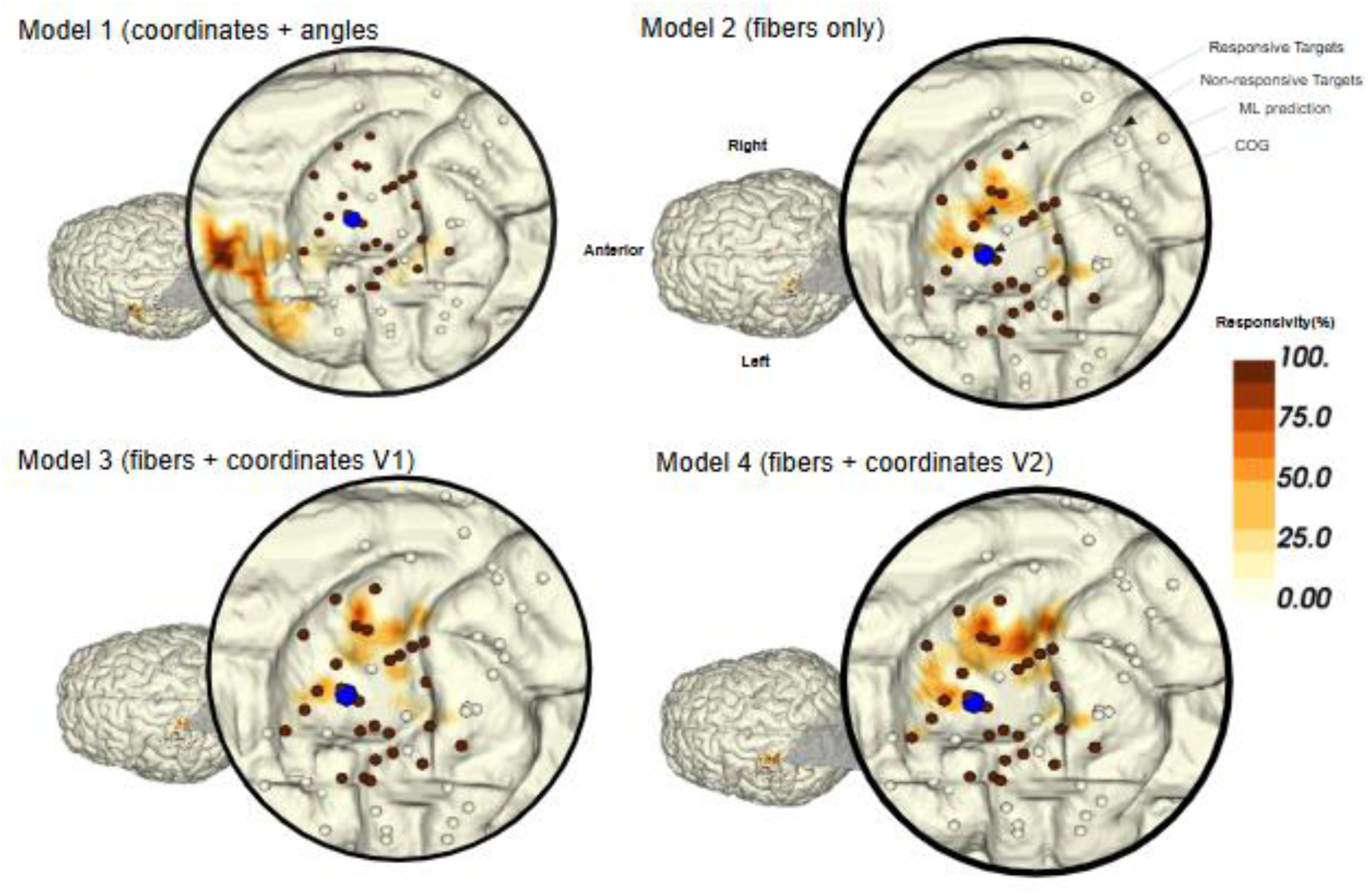
Visualization of predicted and true TMS targets on the cortical surface of Subject 2.

Classification performance was assessed for each subject individually using a 5-fold cross-validation scheme. For each fold, the test accuracy and F1 score were computed. Final performance metrics were then obtained by averaging the results across all five folds.

Model training was conducted on distinct computational platforms according to resource demands. Model 1 (MLP), due to its lower computational complexity, was trained on a local workstation equipped with an Intel® Core™ i7-12700 CPU, 32 GB of RAM, and an NVIDIA GeForce RTX 3060 GPU. In contrast, Models 2 to 4, which involved 3D convolutional architectures, were trained on a dedicated high-performance workstation featuring an AMD Ryzen Threadripper PRO 5975WX processor (32 cores / 64 threads), 128 GB of DDR4 RAM at 3200 MHz, and an NVIDIA RTX A5000 GPU with 32 GB of dedicated memory.

### Responsiveness Brain mapping

The machine learning models were used to predict the responsiveness map of Subject 2, selected due to the motor mapping that encompasses a substantial region of the precentral gyrus and a large area covering the hand cortical area(44). For this subject, a threshold of ≥50 µV was used to classify cortical targets as responsive, with lower values indicating non-responsive regions. These classifications were visualized in a 3D map of the motor cortex, with values ranging from 0 (non-responsive) to 100 (highly responsive).

In this experiment, true experimental data were used to train and test the prediction of target points. To better illustrate the spatial distribution, the models also predicted the responsiveness of the whole cortical area encompassed by the experimental targets. The results are shown as a predicted cortical map whose intensity was interpolated across the primary motor cortex.

## Results

### Machine learning models

Model performance was evaluated in a within-subject setting, where training and testing were performed on data from the same individual using five-fold cross-validation. The data labels used to train the models were defined by splitting the stimulated targets based on the median of their mean MEP amplitudes. This approach was adopted to create two classes of approximately equal size. For most volunteers, the threshold is the median MEP amplitude. However, for Subject 2, the median MEP was below 50 µV, which would have led to the misclassification of targets with biologically negligible responses as “responsive”. To avoid this, a fixed threshold of 50 µV was used instead for this subject. A mean score of laterality index between the subjects was 92 ± 8, indicating strong right-hand dominance.

The classification accuracies and F1 score for each subject and model in this condition are summarized in Table 1 and 2 respectively.

**Table 1:** Cross validation 5 folds subjects (accuracy)

|  | Subject 1 | Subject 2 | Subject 3 | Subject 4 | Subject 5 |
| --- | --- | --- | --- | --- | --- |
| Model 1 (coordinates + angles) | **0.5±0.11** | 0.70±0.13 | 0.63±0.14 | 0.81±0.12 | **0.66±0.10** |
| Model 2 (Fibers only) | 0.38±0.07 | 0.67±0.13 | 0.63±0.10 | 0.84±0.10 | 0.56±0.08 |
| Model 3 (Fibers + coordinates V1) | 0.48±0.13 | 0.53±0.07 | **0.72±0.15** | 0.86±0.10 | 0.50±0.06 |
| Model 4 (Fibers + coordinates V2) | 0.4±0.06 | **0.72±0.14** | 0.63±0.07 | **0.88±0.07** | 0.50±0.11 |

The provided table displays the accuracy results of a 5-fold cross-validation performed on five different subjects, comparing four distinct models. Each cell shows the mean accuracy ± standard deviation.

The table 2 displays the F1 score results of a 5-fold cross-validation performed on five different subjects, comparing four distinct models. Each cell shows the mean F1 ± standard deviation

**Table 2:** Cross validation 5 folds subjects (F1 score)

|  | Subject 1 | Subject 2 | Subject 3 | Subject 4 | Subject 5 |
| --- | --- | --- | --- | --- | --- |
| Model 1 (coordinates + angles) | **0.53±0.10** | 0.63±0.20 | 0.57±0.20 | 0.81±0.13 | **0.66±0.09** |
| Model 2 (Fibers only) | 0.14±0.18 | 0.49±0.29 | 0.47±0.17 | 0.81±0.10 | 0.18±0.15 |
| Model 3 (Fibers + coordinates V1) | 0.00±0.00 | 0.00±0.00 | **0.61±0.22** | 0.79±0.21 | 0.29±0.17 |
| Model 4 (Fibers + coordinates V2) | 0.0±0.00 | **0.63±0.22** | 0.46±0.13 | **0.83±0.14** | 0.30±0.16 |

The best-performing subjects were Subjects 2 to 4, who achieved higher F1 scores and accuracy with the hybrid models that combined coil coordinates and fiber information.

### Motor mapping prediction

To demonstrate the practical applicability of our proposed methodology, Figure 6 illustrates how machine learning predictions can be spatially integrated with true stimulation responses on a subject cortical surface; this map corresponds to Subject 2, using predictions generated by four models. This approach enables the automated identification of responsive cortical regions, reducing the need for exhaustive empirical motor mapping procedures. By projecting both predicted and experimentally verified targets onto a 3D anatomical model, we visualize not only the alignment between predicted and actual responsive zones, but also the overall distribution of spatial prediction of APB muscle stimulation probability.

The image below is arranged in a 2×2 grid corresponding to the four machine learning models (each quadrant representing one model), as previously discussed in Figure 4. This figure presents a three-dimensional visualization of predicted and experimentally verified TMS targets overlaid on a cortical surface mesh. The brain model, derived from an anatomical brain segmentation, serves as a reference for the spatial projection of both true and predicted stimulation coordinates. On the left for each quadrant, a global view of the brain is displayed with a color-coded responsibility map, where shades range from light yellow to dark brown according to predicted responsibility amplitude values normalized to a 0–100% scale. These predicted values were obtained from machine learning models trained to estimate cortical APB responsiveness. The right side of each quadrant provides a zoomed-in view of the motor hotspot region, highlighting the spatial correspondence between predicted and actual stimulation responses. True stimulation targets, obtained from experimental sessions and classified based on MEPs, are rendered as spheres. For this subject, a specific classification criterion was applied: each stimulation site was tested with 10 consecutive pulses, and the average peak-to-peak MEP amplitude was computed. If the mean amplitude was below 50 µV, the target was considered non-responsive and is shown as a white sphere; if it exceeded this threshold, it was considered responsive and is rendered in brown. This threshold-based labeling allows for a clear distinction between physiologically active and inactive regions. Predicted stimulation points on the cortical surface were interpolated using a Gaussian kernel, resulting in continuous painted regions across the brain mesh that reflect the predicted level of responsiveness. These regions, inferred by the neural networks, follow the same colormap used for the background interpolation. To enhance visual clarity, only the top 20% of predicted values—based on normalized responsivity predictions—were retained and rendered. Additionally, a blue sphere marks the center of gravity (CoG) of the empirically defined hotspots. This CoG was computed as a weighted average of the most responsive stimulation points, with weights derived from the corresponding MEP amplitudes. This integrative visualization highlights the model’s capacity to approximate physiologically meaningful patterns of cortical excitability. This predictive process can be performed in real time during neuronavigation procedures using Invesalius Navigator plugin.

The responsivity map overlaid on the brain surface uses a color scale from light yellow to dark brown, indicating the predicted likelihood of responsiveness by the machine learning model. By applying the methodology of displaying only the top 20% of predictions (highlighted area), it is possible to restrict the analysis to regions with the highest predicted responsiveness. This approach effectively highlights the main cortical area where the model predicts the greatest excitability, simplifying the interpretation and identification of the primary motor hotspot.

It can be observed that the CoG, marked by the blue sphere, is located slightly away from the region of highest responsive prediction in Models 2–4, and considerably farther in Model 1. This suggests that the most intense physiological response, as predicted by the AI models, may lie outside the geometric center of the experimentally measured responses.

## Discussion

### Machine learning models

The inter-subject variability, the differences in anatomical organization, tractography-derived features. Together, these results highlight the importance of personalized modeling strategies in the classification of responsive cortical targets.

Although the overall classification accuracy already suggests notable differences between subjects, for some individuals, the per-class accuracy diverges considerably. This imbalance indicates that the criterion adopted to label targets as responsive or non-responsive may not have been universally appropriate across all participants.

Our approach involves classifying stimulation targets as responsive or non-responsive using a machine learning model. Determining these classes is crucial as it directly defines the targets that the model should classify. In contrast to previous works, (45), a target was labeled as responsive if the MEP amplitude exceeded 20 times the EMG baseline standard deviation, with individual thresholds defined per subject to account for baseline variability and responses were considered valid only if the amplitude remained below 10 mV and the latency fell between 5–50 ms. In (46) , a site was considered active if at least two out of six stimuli evoked MEPs ≥ 50 µV under resting conditions, or if at least two clearly discernible from background EMG were elicited (active conditions)

Further emphasizing the variability in classification approaches, (47) adopted a fixed threshold, considering any site with a mean MEP amplitude below 0.05 mV as non-responsive, setting those MEPs to zero to exclude background noise. Similarly, (48) classified a site as active if at least two of four stimuli produced discernible MEPs, and the mean amplitude was calculated as the average across all four pulses. These examples illustrate the lack of consensus in the field, highlighting how different methodological choices can influence the interpretation of motor maps.

To demonstrate the importance of how responsive and unresponsive targets are defined, the comparisons presented in the supplementary materials (subchapter Comparisons Between Responsiveness Methods) are provided. detailed comparison between two distinct methods for classifying cortical target responsiveness based on MEP amplitudes in a single subject. For Subjects 1, 3, and 5, a high level of disagreement was observed between the median-based criterion and the fixed 50 µV threshold. In all three cases, more than 48% of the targets below the median were classified as responsive using the 50 µV criterion. This discrepancy likely introduced a class imbalance in the dataset, resulting in a larger proportion of targets being labeled as responsive regardless of their relative amplitude. Such imbalance can hinder the model’s learning process, which aligns with the lower accuracies observed for these subjects, such as Subject 1, whose best model reached only 0.50 ± 0.11.

In contrast, Subject 4 showed a much lower degree of disagreement between criteria: only 25% of the targets below the median were labeled as responsive, and none above the median were misclassified as non-responsive. This more consistent agreement between classification methods led to a more balanced class distribution, which likely contributed to the notably higher model performance for this subject — reaching 0.88 ± 0.07 with Model 4, the highest accuracy across all subject For Subject 2 displays a unique and opposite pattern of disagreement. For this individual, 25.93% of the targets with average MEPs above the median were actually classified as “Non-Responsive” according to the 50 µV criterion. This suggests that, in this case, the 50 µV threshold acts as a stricter or more conservative classifier compared to the median-based method. This stricter classification may result in a cleaner dataset and a more distinct separation between “Responsive” and “Non-Responsive” classes. This increased clarity likely made the task easier for the predictive models and may explain the consistently higher accuracies observed for Subject 2, including values as high as 0.72 ± 0.14 with Model 4.

However, the definition of “responsiveness” remains a key challenge. In this study, responsiveness was determined using MEP amplitude thresholds, but further investigation into this criterion is necessary. It is important to explore other factors that could influence the responsiveness of cortical targets, such as stimulation intensity or temporal characteristics. Additionally, the current methodology could be improved by refining the criteria for defining responsiveness, which would help enhance the accuracy of predictive models and increase the agreement between predicted and observed responses.

### Motor mapping prediction

In Models 2, 3, and 4, the predicted activation patterns shown in Figure 2 were quite similar, despite slight variations in extent and localization. This suggests that even when combining coordinates with fiber information in Models 3 and 4, there is little added value to the predicted activation maps. Both Models 3 and 4 closely resemble Model 2, which only incorporates fiber information, indicating that adding coordinate and fiber data doesn’t significantly change the predicted patterns.

In contrast, Model 1, which uses only coordinates and angles, shows a distinct prediction. It deviates substantially from the other models, showing activation in the premotor area instead of the motor cortex. This suggests that the simpler model, relying solely on spatial coordinates and angles, struggles to accurately predict the motor cortical response, potentially due to a lack of sufficient data for the model to capture the complex structure of cortical excitability. Future work should focus on expanding the sample size and including more diverse populations to validate the robustness of the approach.

## Conclusion

Hotspot hunting is a time-consuming and operator-dependent procedure. Therefore, developing tools capable of automating the visualization and classification of cortical targets holds significant clinical and research value. In this study, motor mapping was performed in five subjects, and the resulting stimulation targets—represented as fiber bundles—were used to train and evaluate four distinct machine learning models. The results demonstrate the strong potential of these models to automatically identify responsive targets, significantly accelerating the motor mapping process..

Furthermore, while the current method focuses on motor mapping, the use of machine learning could be extended to other functional regions of the brain, including sensory and cognitive areas. Integrating additional modalities, such as EEG or fMRI, could allow the model to incorporate functional data, potentially enabling real-time, whole-brain mapping during TMS. This approach could facilitate the automation of functional target identification, offering a more comprehensive understanding of brain networks and enhancing the precision of neuromodulation techniques.

## Title of new section

Declaration of generative AI and AI-assisted technologies in the writing process.

## Statement

During the preparation of this work the author(s) used Google Gemini and ChatGPT to assist with improving textual cohesion. After using this tool/service, the author(s) reviewed and edited the content as needed and take(s) full responsibility for the content of the published article.

## Declaration of Interest Statement

RHM has received consulting fees from Nexstim Plc, unrelated to this study. The other authors declare that they have no known competing financial interests or personal relationships that could have influenced the work reported in this paper.

## CRediT author contribution statement

Lucas dos Santos Betioli: **Conceptualization**, **Methodology**, **Software**, **Investigation**, **Writing, Data Curation – Original Draft.** Renan Hiroshi Matsuda: **Conceptualization**, **Methodology**, **Investigation**, **Writing, Software – Review & Editing.** Thais Cunha Marchetti: **Conceptualization, Investigation**, **Data Curation**, **Writing – Review & Editing.** Carlo Rondinoni, Victor Hugo Moraes and Márcio Campos: **Investigation**, **Data Curation**, **Writing – Review & Editing.** Carlos Ernesto Garrido Salmon: **Writing – Review & Editing.** Renato Tinós: **Funding Acquisition**, **Project Administration**, **Writing – Review & Editing.** Oswaldo Baffa Filho: **Funding Acquisition**, **Project Administration**, **Writing – Review & Editing**

## Acknowledgments

The authors would like to thank Hohana Gabriela, Prof. Dr. Dogu Baran Aydogan for insightful discussions on tractography, and Prof. Dr. Marco Garcia from the Federal University of Juiz de Fora for valuable discussions on the motor and biological aspects of TMS and its biophysical implications. We are also grateful to the technicians Adriano Holanda from the Department of Computing and Mathematics – FFCLRP-USP for technical support on computational resources, and Lourenço Rocha and Fernando Torrieri from the Department of Physics – FFCLRP-USP, as well as the MRI team from the Hospital das Clínicas, University of São Paulo Medical School at Ribeirão Preto, for their valuable assistance. Finally, we acknowledge the Graduate Program in Applied Physics to Medicine and Biology (FAMB-USP) for providing the institutional and academic support that made this research possible.

## Funding

This research has received funding from the Conselho Nacional de Desenvolvimento Científico e Tecnológico (CNPq; grant number 131294/2023-7 and 305827/2023-5), Tandem Industry Academia (decision No. 366), the Research, Innovation and Dissemination Center for Neuromathematics (FAPESP; grant number 2013/07699-0, 2024/07674-1, 2025/06042-4, 2024/03703-7, 2025/07274-6), and CAPES.

