## Supplementary Materials for "Tractography-based transcranial magnetic stimulation prediction using machine learning"

Affiliations:

Equal contribution in first authorship (#)

Corresponding author:

*Lucas dos Santos Betioli

### Supplementary Materials

#### Comparisons Between Responsiveness Methods

Figures 1 to 5, each corresponding to a different subject, illustrate the comparison between two TMS target responsiveness methods. Each data point along the x-axis represents a cortical target, the y-axis displays the mean MEP amplitude in microvolts (μV) from 10 stimuli , while the error bars represent the standard deviation of the MEPs for each target, highlighting the variability in response.

The left panel, titled "Median-Split Method," presents the first classification approach. In this method, a single reference threshold is established by calculating the median of all mean MEP amplitudes across all targets. This threshold is depicted by the horizontal dashed line. Targets with a mean MEP amplitude above this median threshold are classified as "Responsive" and are colored green. Conversely, targets with a mean MEP below the threshold are classified as "Non-Responsive" and are colored red.

The right panel, titled "50 μV Count Method," displays the second classification approach. Here, a target is deemed "Responsive" (colored blue) if more than half of the individual stimulus pulses delivered to it (i.e., more than 5 out of 10 pulses) elicited an MEP with an amplitude greater than 50 μV. If this condition is not met, the target is classified as "Non-Responsive" (colored orange).

Crucially, the right panel also serves to highlight the discordance between the two classification methods. The symbols 'X' and '+' are used to flag targets where the two methods disagree, using the median-split threshold from the first panel as a reference for comparison:

- An 'X' marker indicates a target that was classified as Responsive by the 50 μV count method, but its mean MEP amplitude is actually below the median-split reference threshold.
- A '+' marker indicates a target that was classified as Non-Responsive by the 50 μV count method, yet its mean MEP amplitude is above the median-split reference threshold.

The analysis of the divergence between classification methods ("Median-Split" vs. "50 μV Count") provides a plausible explanation for the variability in model performance.

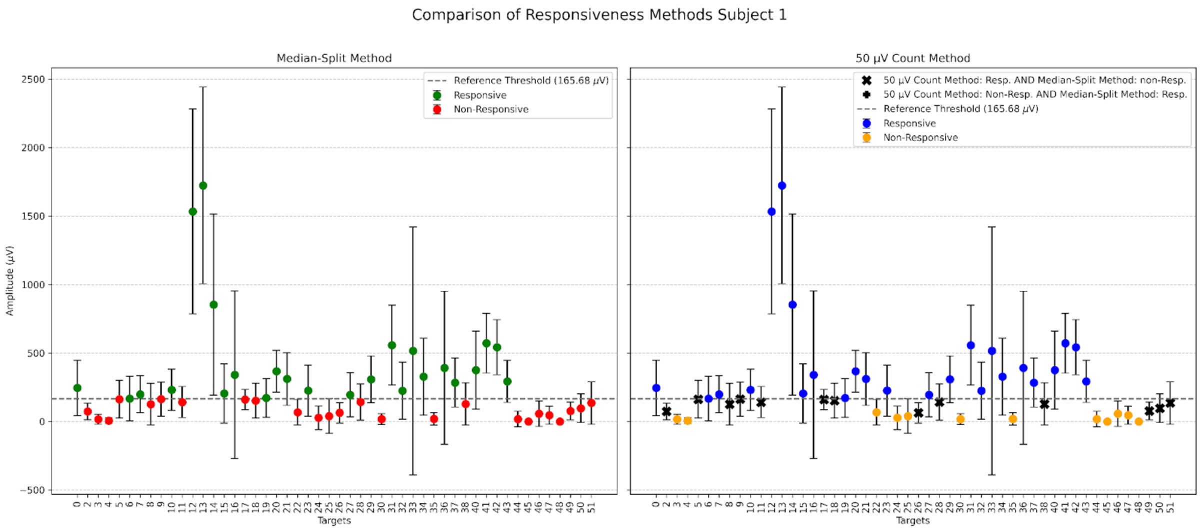


Fig 1: Comparison histogram for Subject 1.

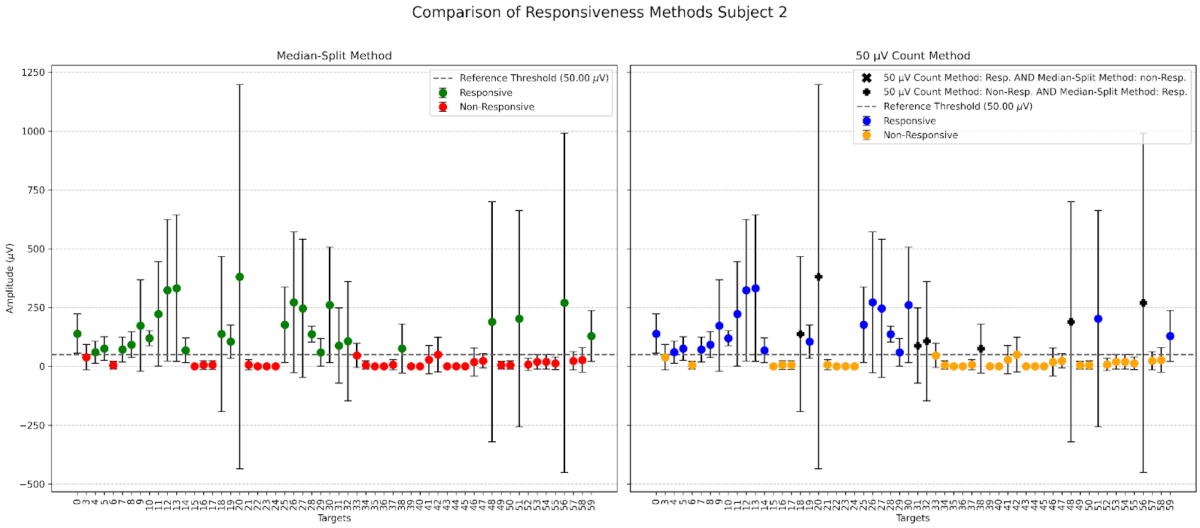

Fig 2: Comparison histogram for Subject 2.

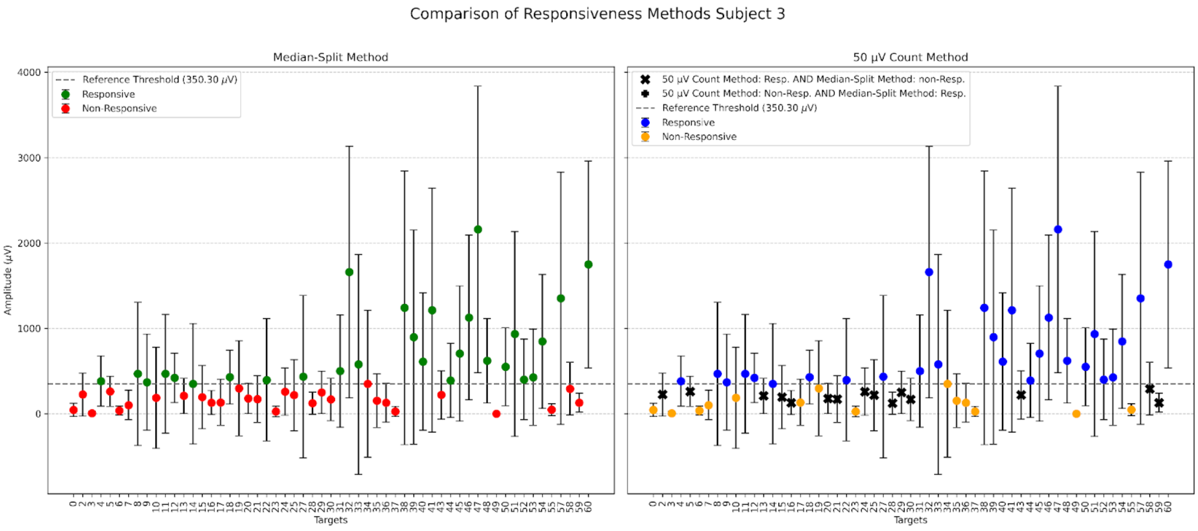


Fig 3: Comparison histogram for Subject 3.

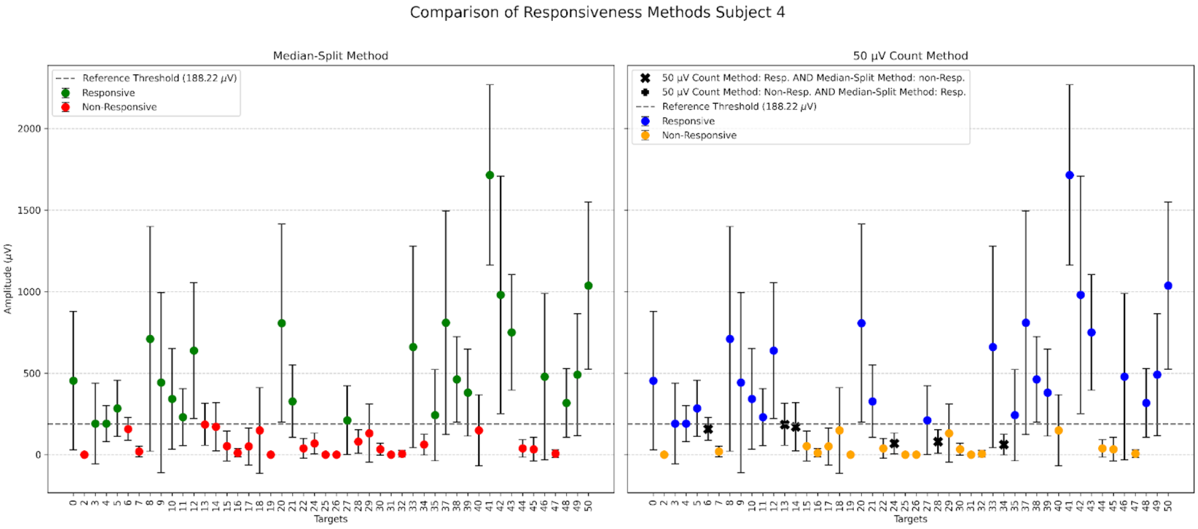


Fig 4: Comparison histogram for Subject 4.

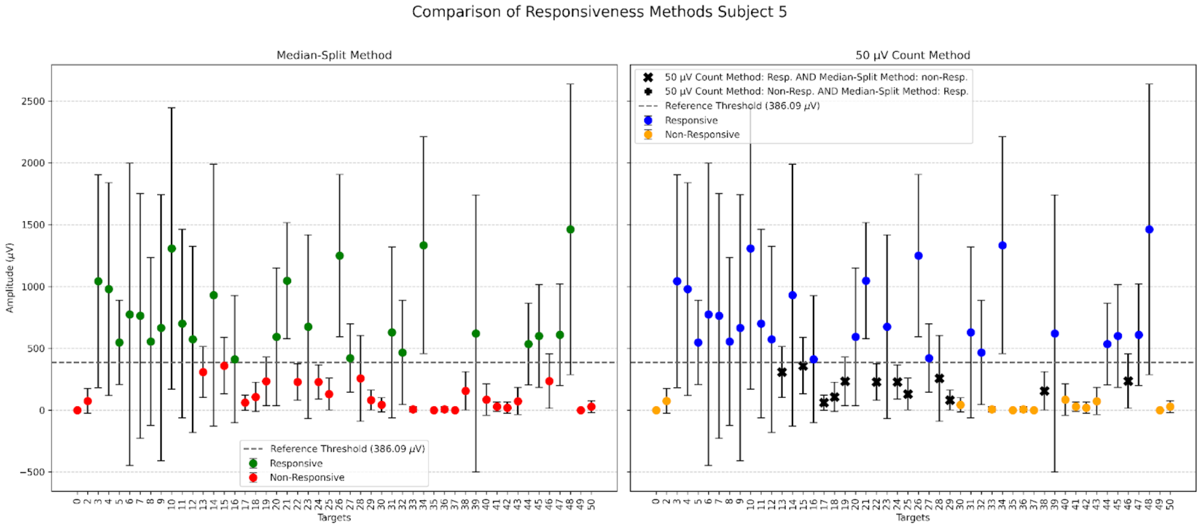


Fig 5: Comparison histogram for Subject 5.

#### Projection of Stimulation Targets onto the White Matter Surface

For each stimulation target selected from the previously defined spatial grid, EMG signals were recorded, and the coil orientation and position—expressed as Euler angles (α, β, γ) and Cartesian coordinates (X, Y, Z)—were represented as a single transformation matrix. Each target projected to white matter and after projection was made the seed tractography This matrix was applied to place a VTK sphere, with a 1.5-unit radius, at each stimulation site. The recon-all FreeSurfer white-matter mesh (exported as an STL file) served as the reference surface: spheres were translated along their local Z-axis in 1-unit increments until the first intersection with the mesh. If no collision occurred within 35 steps, a rigid iterative closest point algorithm (VTK's ICP, up to 1000 iterations) refined the sphere's alignment, ensuring sub-unit precision. The resulting collision coordinates on the white-matter surface were recorded and exported as tractography seed points.
